# The extracellular matrix phenome across species

**DOI:** 10.1101/2020.03.06.980169

**Authors:** Cyril Statzer, Collin Y. Ewald

## Abstract

Extracellular matrices are essential for cellular and organismal function. Recent genome-wide and phenome-wide association studies started to reveal a broad spectrum of phenotypes associated with genetic variants. However, the phenome or spectrum of all phenotypes associated with genetic variants in extracellular matrix genes is unknown. Here, we analyzed over two million recorded genotype-to-phenotype relationships across hundreds of species to define their extracellular matrix phenomes. By using previously defined matrisomes of humans, mice, zebrafish, *Drosophila*, and *C. elegans*, we found that the extracellular matrix phenome comprises of 3-10% of the entire phenome. Collagens (*COL1A1, COL2A1*) and fibrillin (*FBN1*) are each associated with more than 150 distinct phenotypes in humans, whereas collagen *COL4A1*, Wnt- and sonic hedgehog (shh) signaling are predominantly associated in other species. We determined the phenotypic fingerprints of matrisome genes and found that *MSTN, CTSD, LAMB2, HSPG2*, and *COL11A2 and their corresponding* orthologues have the most phenotypes across species. Out of the 42’558 unique matrisome genotype-to-phenotype relationships across the five species with defined matrisomes, we have constructed interaction networks to identify the underlying molecular components connecting with orthologues phenotypes and with novel phenotypes. Thus, our networks provide a framework to predict unassessed phenotypes and their potential underlying molecular interactions. These frameworks inform on matrisome genotype-to-phenotype relationships and potentially provide a sophisticated choice of biological model system to study human phenotypes and diseases.

**Highlights:**

- 7.6% of the human phenome originates from variations in matrisome genes
- 11’671 phenotypes are linked to matrisome genes of humans, mice, zebrafish, *Drosophila*, and *C. elegans*
- Expected top ECM phenotypes are developmental, morphological and structural phenotypes
- Nonobvious top ECM phenotypes include immune system, stress resilience, and age-related phenotypes

## Introduction

A principle goal in biology is to link molecular mechanisms and biological pathways to phenotypes. Recent systems-level approaches of genome-wide association studies (GWAS) and phenome-wide association studies (PheWAS) have been harnessed to generate and map genotype-to-phenotype relationships. These efforts are invaluable for clinical research spearheaded by linking electronic health records to medically relevant PheWAS, thereby associating genetic variants to human phenotypes, pathologies, and diseases [1–4]. Assuming that conserved genes and pathways result in comparable phenotypes across species, knowing the genotype-to-phenotype relationship in one species could inform on the existence of a similar genotype-to-phenotype relationship in another species. The phenome is the entire set of phenotypes resulting from all possible genetic variations [5,6]. In order to make trans-species genome-phenome inferences, the phenomes of several species need to be known. The human phenome is far from reaching saturation [6]. We estimate that the current human phenome consists of 7’037 unique phenotypes (Source: Database Monarch Initiative; timestamp 23.08.2019). From the entire human phenome, several sub-phenomes have been characterized and defined, including the behavioral phenome [7], the aging phenome [8,9], and the disease phenome [10–12]. These genotype-to-phenotype relationships can be used to build networks and pathways.

In particular, the human disease phenome consists of over 80’000 genetic variants associated with human diseases, of which cardiovascular diseases and skin and connective diseases share the most pathways [12]. These diseases are enriched in variants closely located or found in collagens or extracellular matrix genes. Cells secrete proteins, such as collagens, glycoproteins, and proteoglycans that integrate and form matrices in the extracellular space. The extracellular matrix (ECM) provides structural support, is important for cell-cell communication, and cellular homeostasis [13,14]. The ECM has emerged as a key feature for overall health or disease, with several clinical trials targeting to modify the ECM [15–18]. Furthermore, recent approaches that combine PheWAS from mice to human or zebrafish to human revealed clinically relevant variants in collagen *Col6a5* or in *Ric1* important for collagen-secretion [19,20]. This indicates that a phenome-based approach across species can be of translational value. However, a defined phenome of collagens and other extracellular matrix genes is currently missing for any species.

Here, we mine publicly available databases to establish the phenome of extracellular matrices across species. We take advantage of the large collection of over two million genotype-to-phenotype associations that integrates data across hundreds of species from more than 30 different curated databases provided by the Monarch Initiative [21]. For the ECM phenome we include all genes that assemble the matrisome. The matrisome is the compendium of all possible gene products that either form, remodel, or associate with the extracellular matrix [22]. The matrisome of humans consists of 1027 proteins [23], for mice 1110 proteins [23], for zebrafish 1002 proteins [24], for *Drosophila* 641 proteins [25], and for *C. elegans* 719 proteins [26]. This corresponds to roughly 4% of their genomes are dedicated to ECM genes. We find a similar contribution of 3-10% (overall 6.7 %) of the phenome consisting of 42’558 ECM-specific gene-to-phenotype associations. Across these five species, 639’431 gene-phenotype associations were recorded consisting of 34’818 distinct phenotypes. We identify the phenotypic landscape of the matrisome. Our genotype-to-phenotype interaction maps for matrisome genes reveal unstudied interaction when compared across species.

## Methods and Materials

### Matrisome phenotype-to-genotype relationships

The phenotype-gene association table was obtained from the Monarch Initiative (Source: Database Monarch Initiative; timestamp 23.08.2019) [21]. This phenotype-gene association table was modified to remove duplicated species-gene-phenotype relationships (< 0.002% in total) and unspecified organisms were removed (Supplementary Table 1). The genotype-to-phenotype information was then combined with the defined matrisomes of five species: the matrisome of humans [23], mice [23], zebrafish [24], *Drosophila* [25], and for *C. elegans* [26]). To enable cross-species comparisons among the species with defined and undefined matrisomes, we mapped all genes documented in the Monarch Initiative database to their corresponding orthologues. The homology mapping was achieved by using the homologene database by the National Center for Biotechnology Information (NCBI) implemented in the homologene R package (Ogan Mancarci (2019). homologene: Quick Access to Homologene and Gene Annotation Updates. R package version 1.4.68.19.3.27. https://CRAN.R-project.org/package=homologene). The human homology information subset and its associated phenome was then further analyzed. All data analysis was performed using the purrr and dplyr R packages (Lionel Henry and Hadley Wickham (2019). purrr: Functional Programming Tools. R package version 0.3.3. https://CRAN.R-project.org/package=purrr; Hadley Wickham, Romain François, Lionel Henry and Kirill Müller (2019). dplyr: A Grammar of Data Manipulation. R package version 0.8.3. https://CRAN.R-project.org/package=dplyr).

### Grouping of orthologous phenotypes

To facilitate the comparison between species, the phenotypes were simplified and grouped to form larger phenotype collections (Supplementary Table 2). These phenotype collections enable both cross-species comparisons by avoiding species-specific terminology, as well as, generating a broader overview of the processes involved. Substring matching was used to assimilate similar phenotypes to phenotype groups, while assuring that each original phenotype only belonged to a single phenotype group using the stringr R package (Hadley Wickham (2019). stringr: Simple, Consistent Wrappers for Common String Operations. R package version 1.4.0. https://CRAN.R-project.org/package=stringr). Capitalization and terminal white spaces were ignored.

### Data analysis and visualization

The circular dendrograms were generated with a single hub gene as central node connected to the species in which it was observed and the phenotypes as terminal nodes. The interaction network for each collection of gene-phenotype interactions was computed by extracting the genes, which are associated with the largest number of phenotypes followed by selecting the phenotypes, which comprise the largest number of genes. The edges of the generated network were then facetted by species and their weight set according to the number of species in which a phenotype was observed for the orthologues of a particular gene. Network analysis and visualization was performed using the igraph and ggraph R packages (Csardi G, Nepusz T: The igraph software package for complex network research, InterJournal, Complex Systems 1695. 2006. http://igraph.org; Thomas Lin Pedersen (2018). ggraph: An Implementation of Grammar of Graphics for Graphs and Networks. R package version 1.0.2. https://CRAN.R-project.org/package=ggraph). In addition, the R packages ggplot2, ggforce and ggpubr were used (H. Wickham. ggplot2: Elegant Graphics for Data Analysis. Springer-Verlag New York, 2016; Thomas Lin Pedersen (2019). ggforce: Accelerating ‘ggplot2’. R package version 0.3.1. https://CRAN.R-project.org/package=ggforce; Alboukadel Kassambara (2019). ggpubr: ‘ggplot2’ Based Publication Ready Plots. R package version 0.2.2. https://CRAN.R-project.org/package=ggpubr). Illustrations were generated using Inkscape version 0.92.

## Results

### The matrisome-phenome across species

To define the extracellular matrix phenome, we sought to identify the overall contribution of matrisome genes to the phenome for each species. In particular, to assess the suitability for this approach, we first plotted the phenotypic landscape in relation to genotypes across species (Fig. 1A). We found 7’037 unique phenotypes comprising the human phenome, from which 3’809 unique genes are linked to these phenotypes (Supplementary Fig. 1). Fortunately, the species with a good phenotype-to-genotype ratio are the ones that have a fully defined and characterized matrisome [23–26]. We used these defined matrisome gene lists to determine their contribution to the phenomes of humans, mice, zebrafish, *Drosophila*, and *C. elegans* (Fig. 1B-1F). The contribution of the matrisome to the human phenome is 7.6%, with a comparable 2.8-9.5% across species (Fig 1B-F). Thus, the matrisome is involved in a number of phenotypes, but only corresponds to a maximum of 10% of the overall recorded phenome across species.

**Fig. 1:**
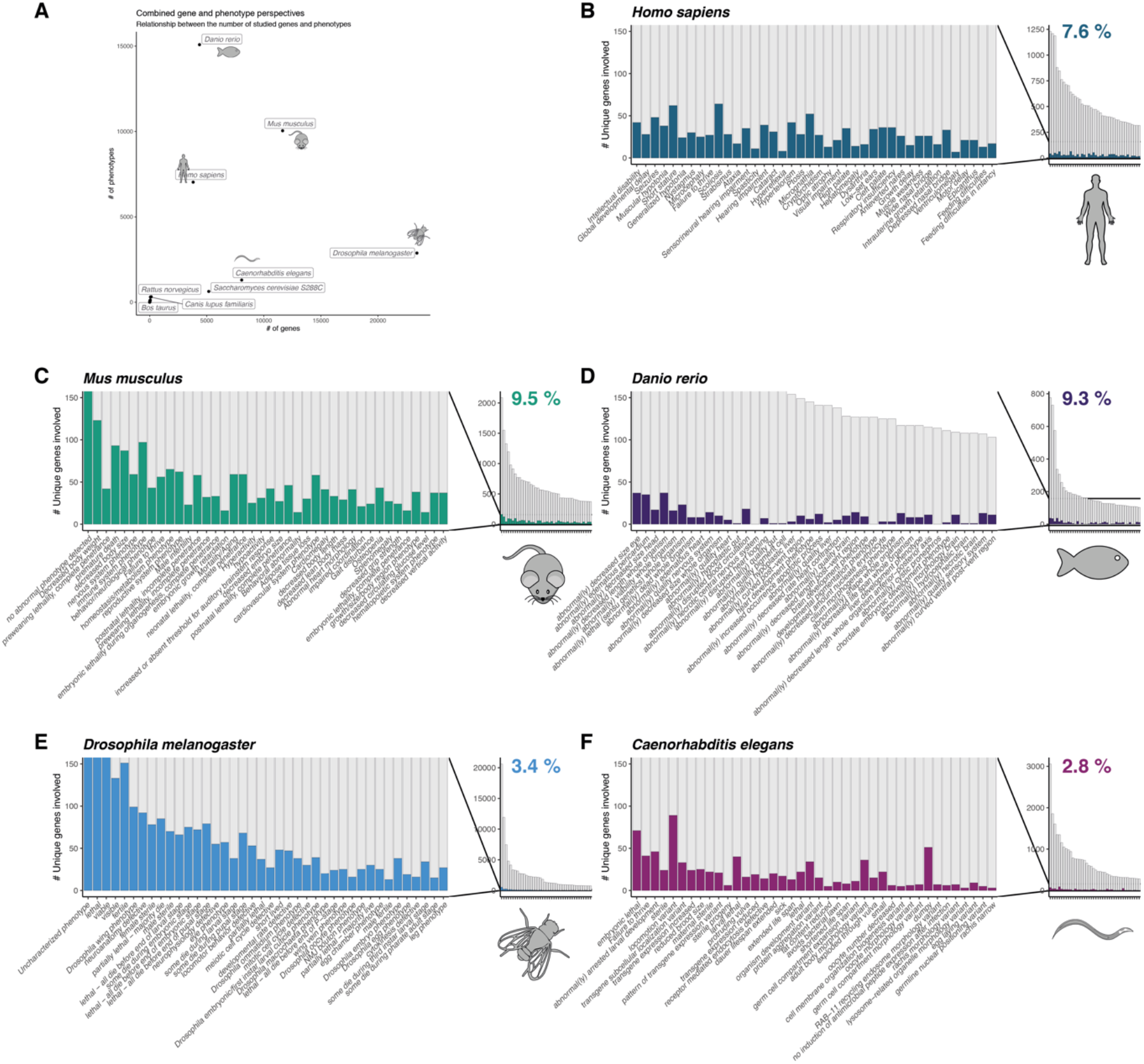
The contribution of the matrisome to the phenome across species. The relationship between the number of genes associated with at least one phenotype and the total number of characterized phenotypes (phenome) linked to at least one gene are shown across species (A). The species with defined matrisomes are depicted in separate panels with the number of genes associated with each phenotype shown on the y-axis and the 40 phenotypes with the highest number of total genes involved are shown on the x-axis for humans (B), for mouse (C), for zebrafish (D), for *Drosophila* (E), and for *C. elegans* (F). The fraction of genes, which are members of each species’ matrisome are highlighted in color. The right panel provides an overview of the gene-to-phenotype distribution, while the left panel highlights the contribution of the matrisome by cropping the y-axis. The fraction of matrisome genes associated with the phenotypes compared to all genes is shown as a percentage.

### The phenotypic signatures of matrisome genes across species

Next, we determined which phenotypes are associated with the greatest number of matrisome genes. For humans the top three phenotypes are “scoliosis”, “short stature”, and “micrognathia” (Supplementary Fig. 2). By contrast, the top three phenotypes for mice are “decreased body weight”, “immune system”, “premature death”, for zebra fish (“decreased eye size”, “decreased length whole organism”, “lethal”), for *Drosophila* (“uncharacterized phenotype”, “viable”, “lethal”), and for *C. elegans* (“locomotion variant”, “embryonic lethal”, “dumpy”; Supplementary Fig. 2). Given that certain phenotypes are species specific, for instance “dumpy” stands for short and fat *C. elegans*, we realized that we cannot compare the phenotypic landscape across species using this conventional phenotypic nomenclature. To overcome this obstacle, we grouped phenotypes into novel phenotype collections. For instance, the phenotypic category “altered body size” includes phenotypes with the key words: body weight, body size, body length, growth phenotype, dwarf, small, dumpy, stunted, tall and others (Materials and Methods, Supplementary Table 1 and 2). With this approach, we were able to group 86.8% of the total 716’644 species-gene-phenotype associations (Supplementary Table 2). We identified sterility, development, and altered body size as the most dominant phenotypic categories across species, and for instance connective tissue and skin phenotypes for vertebrate and poor viability for invertebrates (Fig. 2). Similarly, besides skin and connective tissue, many morphological phenotypes affecting bone, eye, brain, muscle, and the cardiovascular system are represented in these top categories (Fig. 2). Unexpectedly, aging-related phenotypes, stress resilience, and immune systems phenotypes were ranked across species among these top ECM phenotypic categories (Fig. 2). Since we base our observations on five species, we extended our analysis to incorporate all known genotype-phenotype associations across 44 species. Thereby, we were able to extend the matrisome by homology mapping to three additional species (*Bos taurus* (cattle), *Rattus norvegicus* (rat), *Canis lupus familiaris* (dog) (Supplementary Table 3). Then, we used these matrisome input gene lists to define their matrisome-phenome. We found similar top phenotypes either for the ungrouped or with our comparable phenotypic categories (Supplementary Fig. 3 and 4). Thus, our analysis revealed known ECM phenotypes but also unexpected phenotypes related to the ECM with high penetrance across species.

**Fig. 2:**
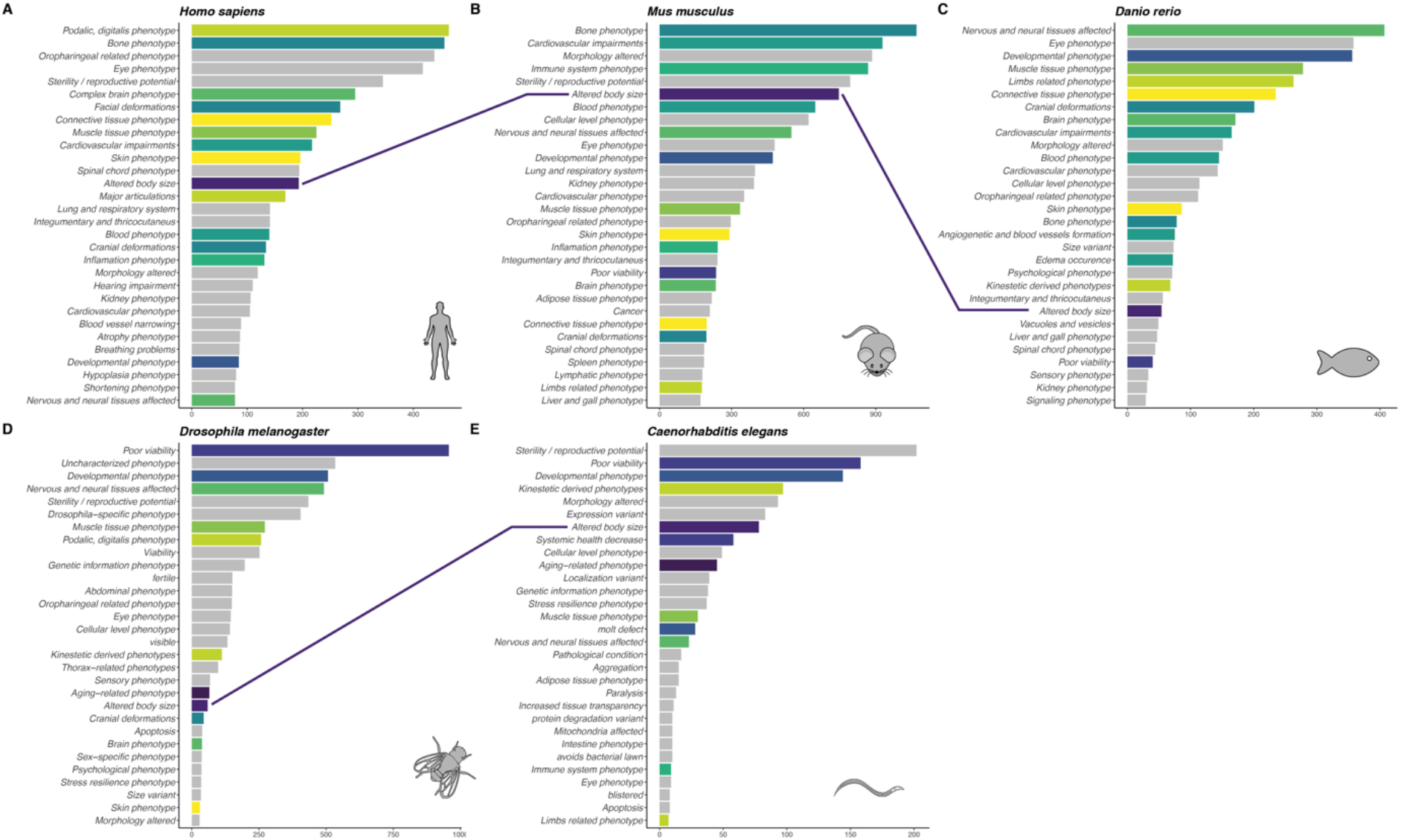
The matrisome-phenome across species. Using the matrisome composition for each species we show the phenome space associated with it for humans (A), mice (B), zebra fish (C), *Drosophila* (D), and *C. elegans* (E). Each species is depicted in a separate panel with the 30 largest phenotype groups on the y-axis and the number of unique gene/phenotype associations contained in each group on the x-axis. Subsets of phenotypes and analog phenotypes across species are color-coded to ease the comparison (i.e., “altered body size” in purple). For illustration we connected the “altered body size” phenotype with a purple line. The phenotypes meaning wear slightly different meaning across species due to different morphologies as i.e. *C. elegans* eye phenotype, which refers to anything related to phototaxis (e.g., UV-light sensing), while the limbs related phenotype captures phenotypes related to the animal’s tail.

### Top ranked-matrisome gene signature of the ECM-phenomes

To understand which genes are the major drivers of multiple distinct phenotypes, we ranked matrisome genes based on their number of associated phenotypes (Fig. 3, Supplementary Fig. 5). For humans, collagens (*COL2A1, COL1A1, COL5A1*), fibrillin (*FBN1*), and growth differentiation factor (*GDF5*) are each associated with more than 150 distinct phenotypes (Fig. 3A). By contrast, for mice, zebrafish, *Drosophila*, and *C. elegans*, Wnt- and sonic hedgehog (shh) signaling and other matrisome-secreted factors are predominantly associated with the greatest number of distinct phenotypes (Fig. 3B-E, Supplementary Fig. 5). Except for zebrafish, the highly conserved collagen type IV (*COL4A1/emb-9*) is associated with range of 20-100 phenotypes across species (Fig. 3). Overall, members of each matrisome category are represented and comparable conserved gene families are linked with numerous phenotypes.

**Fig. 3:**
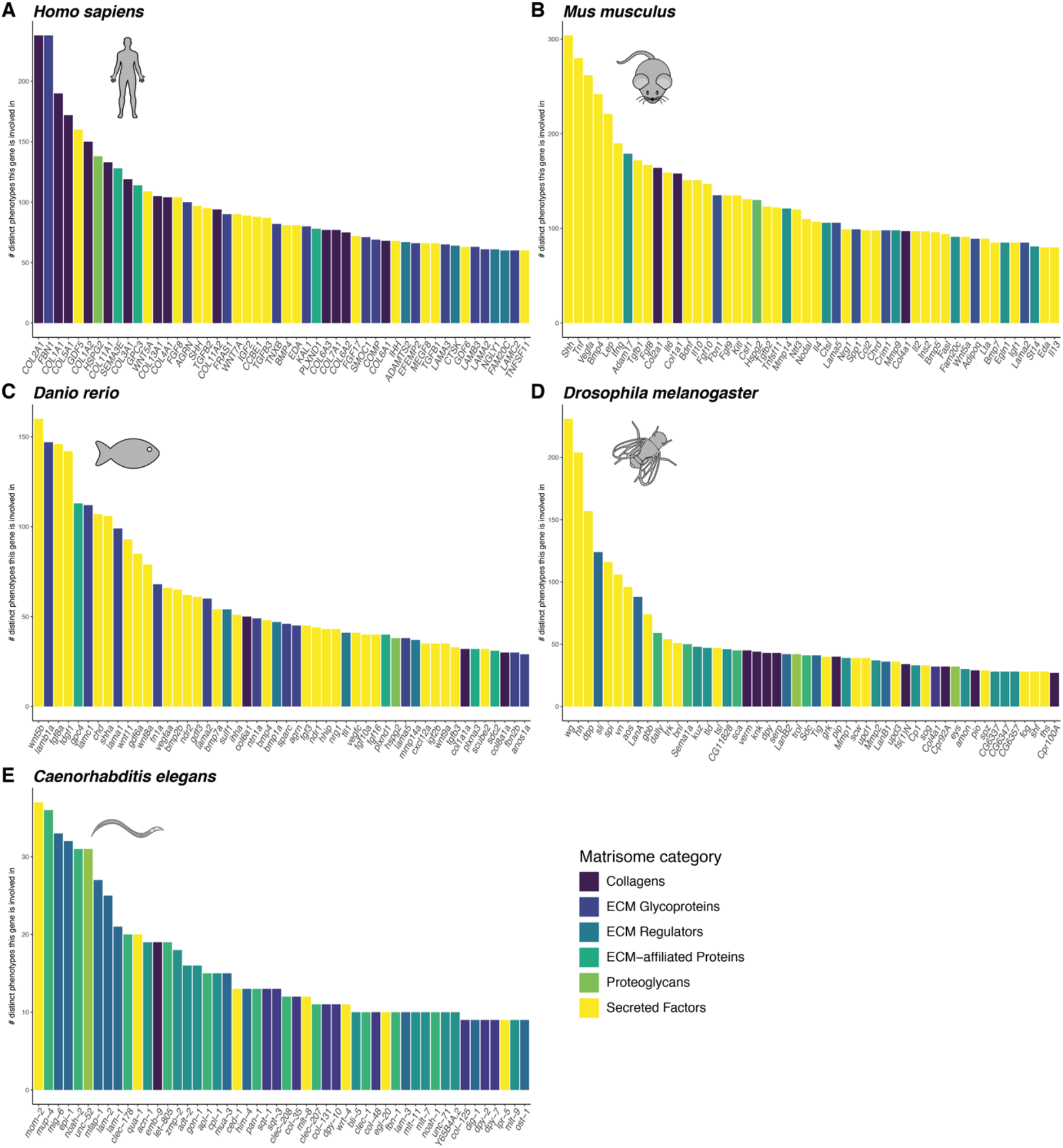
The degree of phenotypic impact of matrisome genes. The number of unique phenotypes linked to individual matrisome gene is shown in decreasing order for the top 50 genes. The gene name is displayed on the x-axis and the number of unique phenotypes on the y-axis. The color of the bars refers to the matrisome category for each gene.

### Phenotypic finger prints associated with matrisome genes

Using our phenotype grouping approach, we observed that certain genes were associated with comparable phenotypes across species. For this comparison, we used our phenotypic categories and plotted all matrisome genes as dendrograms connecting the gene with its phenotypes through the species in which they were observed (Fig. 4, Supplementary Fig. 6). The most phenotypically conserved gene is the secreted muscle growth controlling myostatin (MSTN) protein showing muscle, fat, and body size phenotypes in seven species (Fig. 4A, B). This is not surprising, since either loss or pharmacological inhibition of myostatin leads to almost a doubling of muscle mass, a trait applied in livestock and a potential target for treating muscle wasting diseases [27]. Next is the cathepsin D (CTSD) gene, which is a lysosomal aspartyl protease that can be secreted into the extracellular space to remodel ECM and is involved in many physiological functions, including skin and neuronal development, lipofuscin (age-pigment) removal, and apoptosis [28]. Our analysis of CTSD showed the full phenotypic spectrum across six species with many conserved phenotypes but also species-specific phenotypes (Fig. 4C, Supplementary Fig. 6). Similarly, laminin (LAMB2), heparan sulfate proteoglycan (HSPG2), collagen (COL11A2) showed comparable orthologues phenotypes across multiple species ranging from altered body size to aging-related phenotypes (Fig. 4D-F). The phenotypic finger prints of all matrisome genes is shown in Supplementary Fig. 6 and 7. Thus, we have defined the phenotypic landscape of matrisome genes across species.

**Fig. 4:**
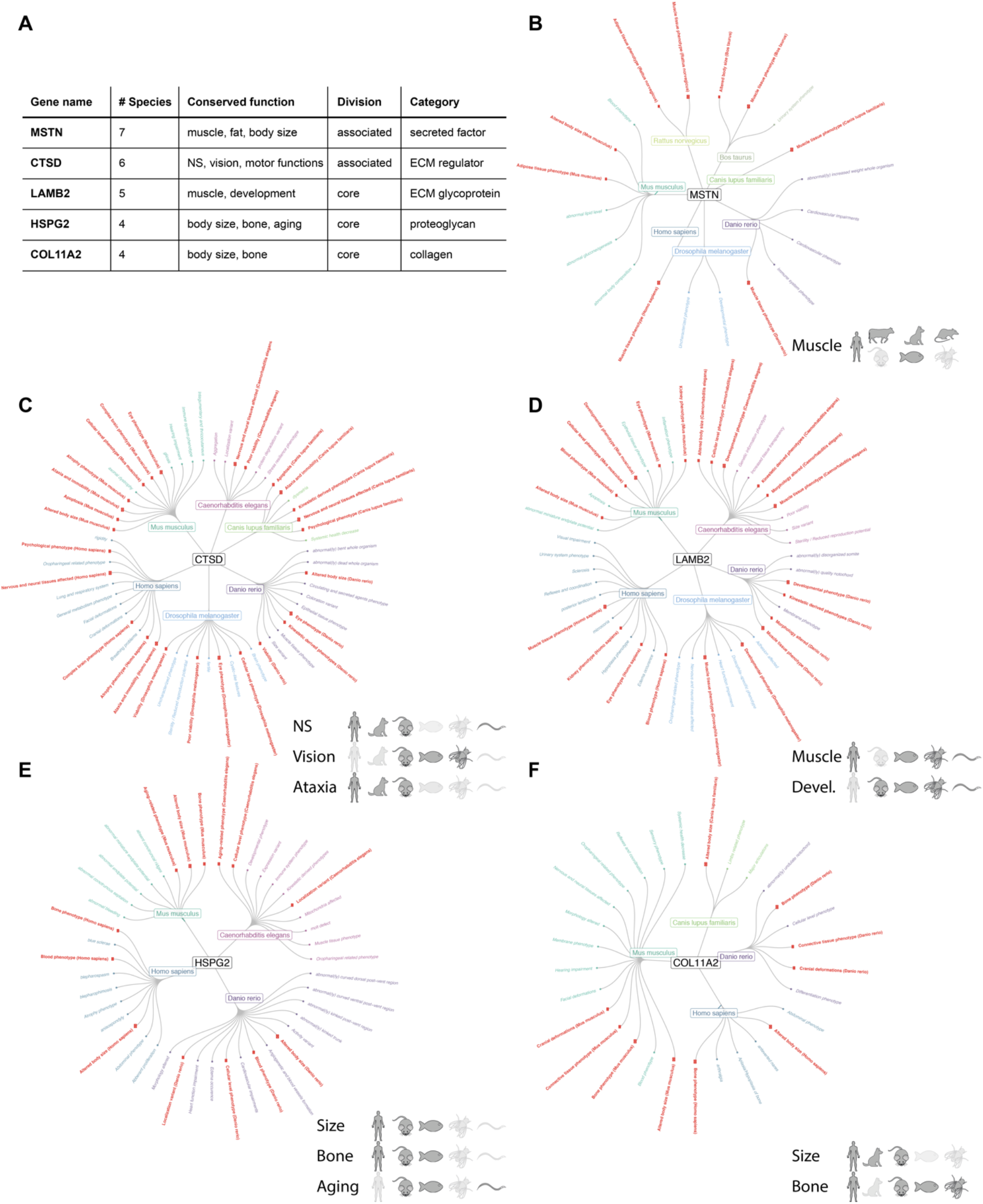
Phenotypic fingerprint of the most conserved matrisome members. The most conserved genes displaying phenotypes across the most species are shown in decreasing order (A). The table displays the human gene name, the number of species presenting a phenotype associated with the matrisome gene, the overall phenotypic signature observed for this gene, as well as the matrisome category and division are shown. Circular dendrograms depict the five most conserved matrisome genes in more detail (B-F). The human orthologue is shown in the middle connected to the species, in which the gene was associated with a phenotype, and then each connected to terminal nodes depicting the grouped phenotypes. Phenotypes that were observed in more than one species are highlighted in red and the size of the terminal node reflects the number of species this phenotype was observed in. At the bottom right of each circular dendrogram panel a graphical summary is provided highlighting the species in which by homology a phenotype group was observed (dark grey) or absent (light grey).

### Network analysis to identify novel phenotypes and underlying molecular mechanisms

We next asked whether we could use this phenotypic landscape of matrisome genes to inform on potential phenotypes not assessed in humans. Furthermore, could we use gene-phenotype pairs in one species to infer novel interactions in another species? In order to achieve this, we built matrisome-genes-to-phenotype networks across species for all matrisome categories and divisions. These networks are shown in Supplementary Fig. 8 and 9. We chose the three core-matrisome categories, which include ECM glycoproteins, collagens, and proteoglycans, to build a network of the five most abundant interspecies gene-to-phenotype associations for *H. sapiens, M. musculus* and *D. rerio* (Fig. 5). We positioned these top five conserved genes and phenotypes for each species at the same location in our graphical network (Fig. 5). We identified the conserved gene-to-phenotypes interactions (bold arrows), but also interactions that only occur in a single species (light arrows; Fig. 5). For instance, genetic variants in neuromuscular junction protein agrin are associated in humans solely with a “sterility” phenotype, whereas in mouse in addition to sterility, agrin is associated with altered body size, morphology, and cardiovascular impairments (Fig. 5). Next, we selected the most connected matrisome genes and mapped the genotype-to-phenotype relationships across species (Fig. 6). Myostatin (*MSTN*) and collagen type VII (*COL7A1*) are the most conserved drivers for muscle or skin phenotypes, respectively (Fig. 6). Laminin (*LAMA2*), heparan sulfate proteoglycan (*HSPG2*), and sonic hedgehog (*SHH*) gene build a strong interaction network among brain, muscle, and cardiovascular phenotypes (Fig. 6). Curiously, only in mice is matrix metallopeptidase 9 (*MMP9*) implicated with these three phenotypes (brain, muscle, and cardiovascular), but also with immune system and proliferation phenotypes (Fig. 6). Taken together, our graphical networks build a framework to predict unassessed phenotypes and their potential underlying molecular interactions.

**Fig. 5:**
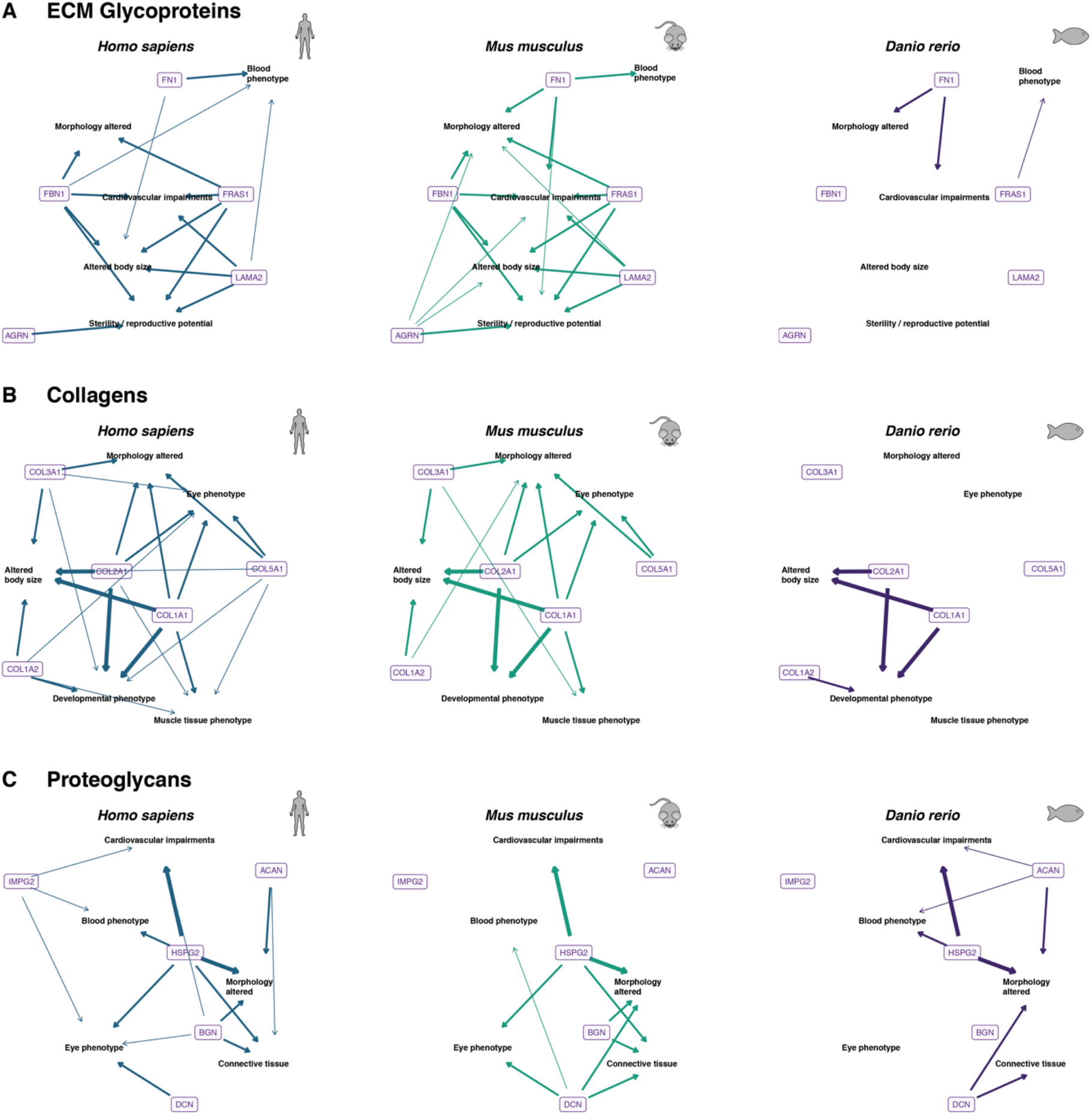
Phenotypic implications of the structural matrisome categories on vertebrates. The matrisome is grouped into multiple categories according to the localization and composition of the extracellular proteins. The members of the ECM glycoprotein category, collagens, and proteoglycans constitute the foundation of the matrisome responsible for the structural integrity of the extracellular matrix. Here the most abundant inter-species gene-to-phenotype associations are highlighted for *H. sapiens, M. musculus* and *D. rerio*. Genes are shown as human orthologues in colored labels while phenotype groups are given as plain text. The arrows connect genes to phenotype groups if the gene has been associated with at least one phenotype belonging to the phenotype group. The width and opacity of the connection reflects the degree of conservation of the gene-to-phenotype association across species.

**Fig. 6:**
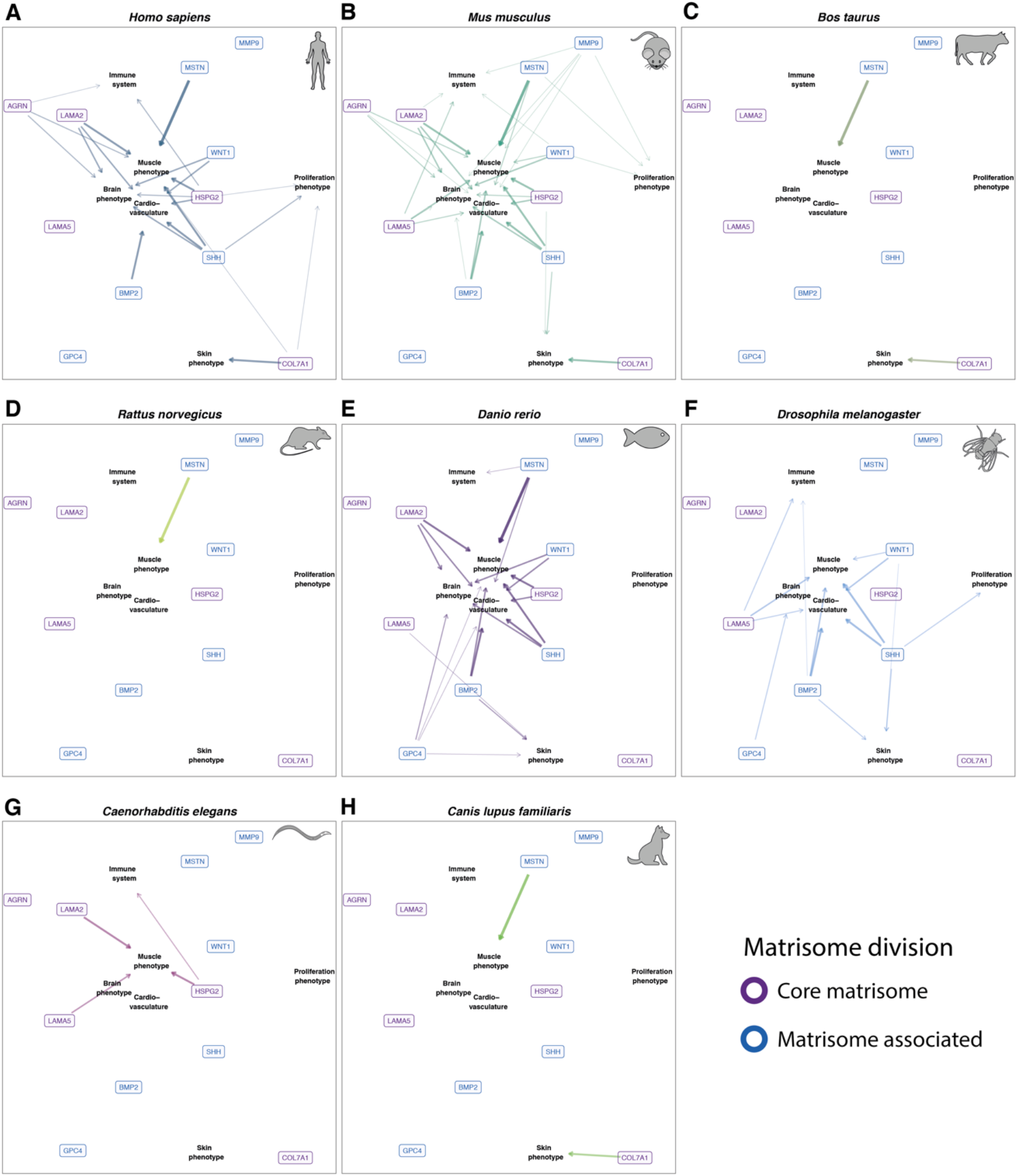
Phenotypic implications of the entire matrisome. A selected subset of the most connected genes of the entire matrisome and phenotypes across species are displayed. Genes are shown as human orthologues color-coded by matrisome division, while phenotype groups are shown as plain text. The arrows connect genes to phenotype groups when the gene is associated with at least one phenotype of the phenotype group. The width and opacity of the connection reflects the degree of conservation of the gene-to-phenotype association across species.

## Figure Legends

**Supplementary Fig. 1:**
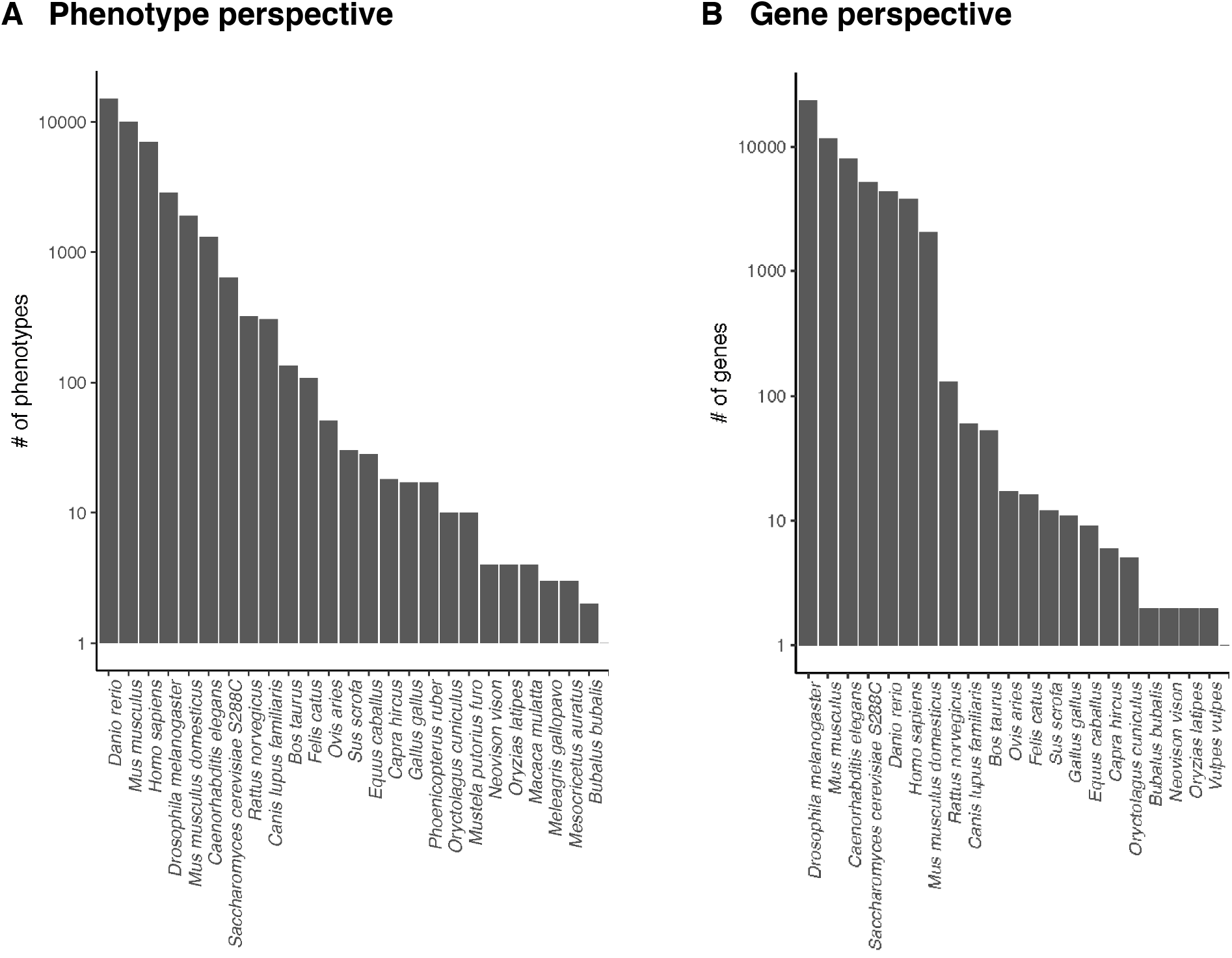
The phenome and number of genes associated with phenotypes. (A) The phenome across species plotted as number of phenotypes. (B) The number of unique genes associated with phenotypes across species.

**Supplementary Fig. 2:**
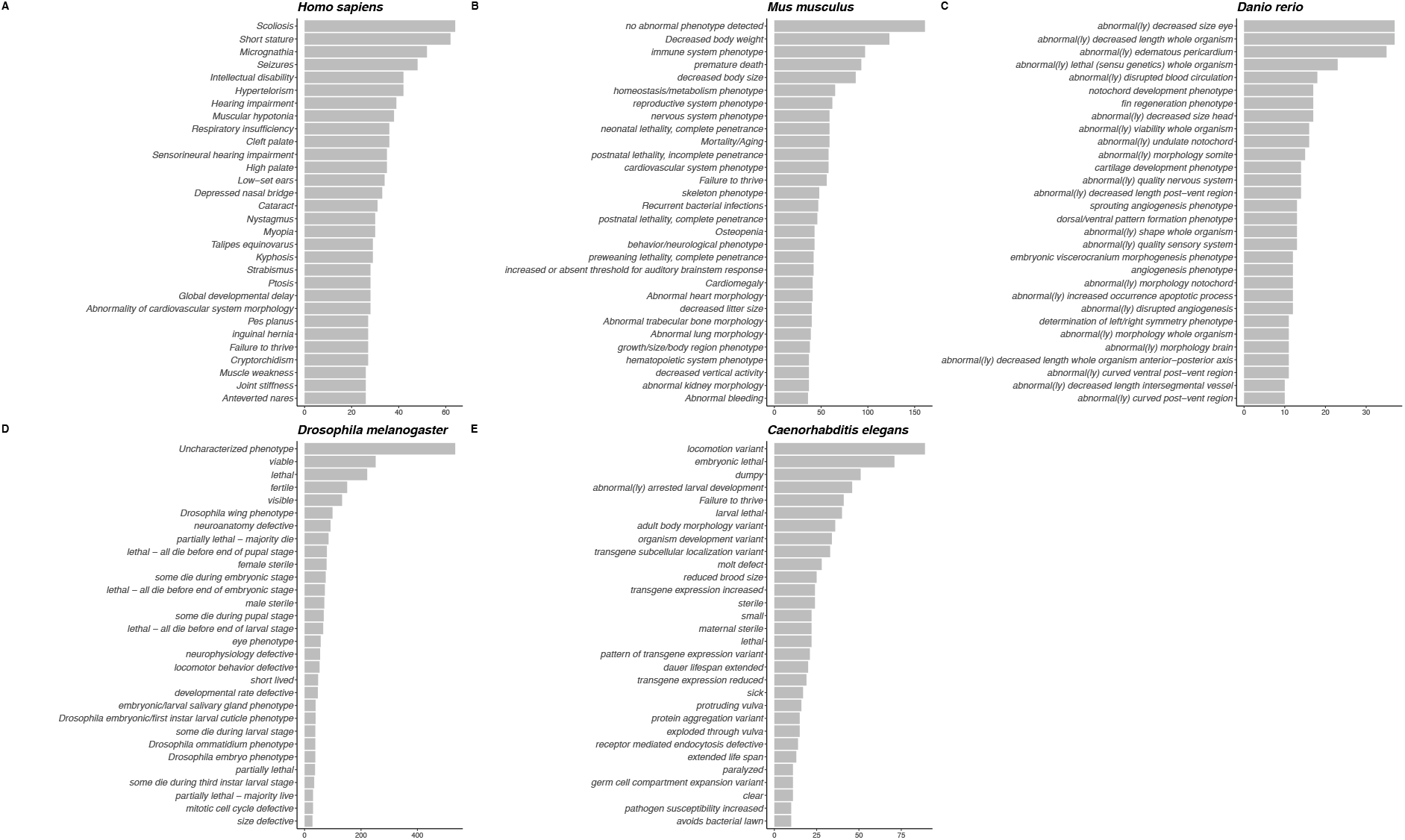
Ungrouped phenotype space across species: Using the matrisome composition for each species and the ungrouped phenotypes, the associated phenome space for humans (A), mice (B), zebra fish (C), *Drosophila* (D), and *C. elegans* (E) are shown. Each species is depicted in a separate panel with the 30 largest phenotype/gene association groups on the y axis and the number of unique gene/phenotype associations contained in each group on the x-axis.

**Supplementary Fig. 3:**
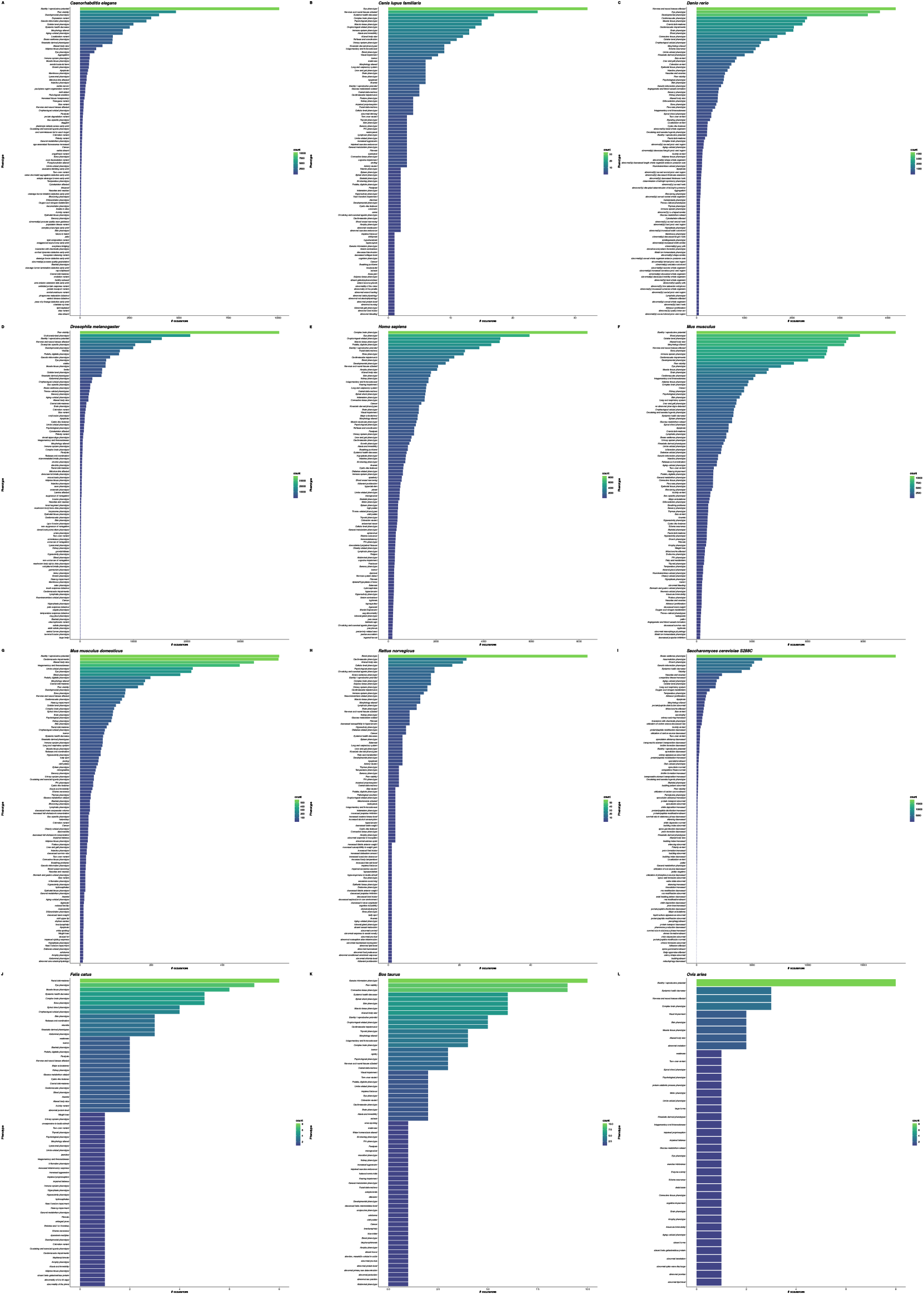
The whole matrisome genome-phenome across species (grouped) All gene/phenotype associations of every species are shown for 44 species. The number of unique gene-to-phenotype associations within every phenotype group are shown arranged in descending order.

**Supplementary Fig. 4:**
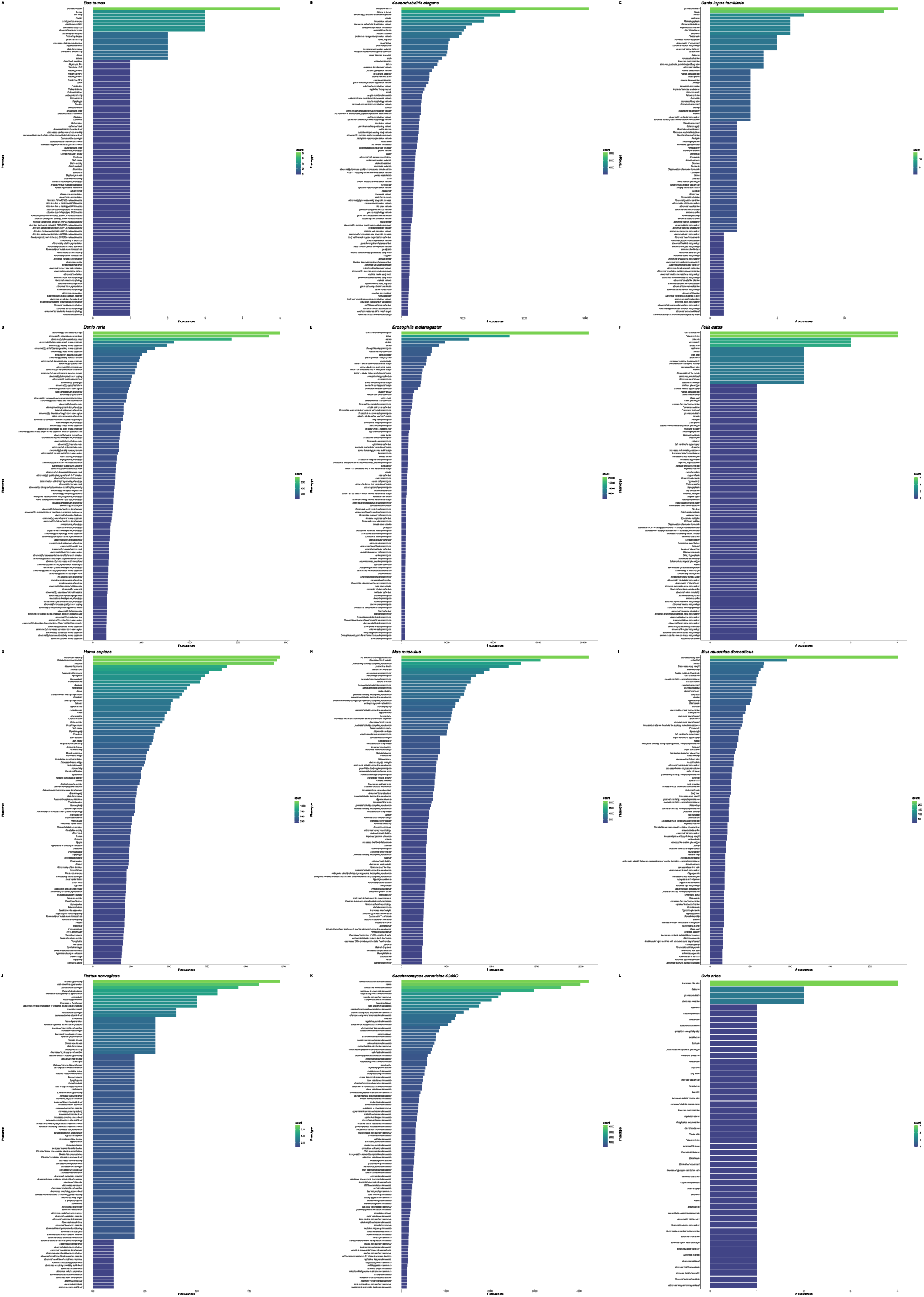
The whole matrisome genome-phenome across species (ungrouped) All gene/phenotype associations of every species are shown for 44 species. The number of unique gene-to-phenotype associations with every phenotype are shown arranged in descending order without assembling phenotypes into groups.

**Supplementary Fig. 5:**
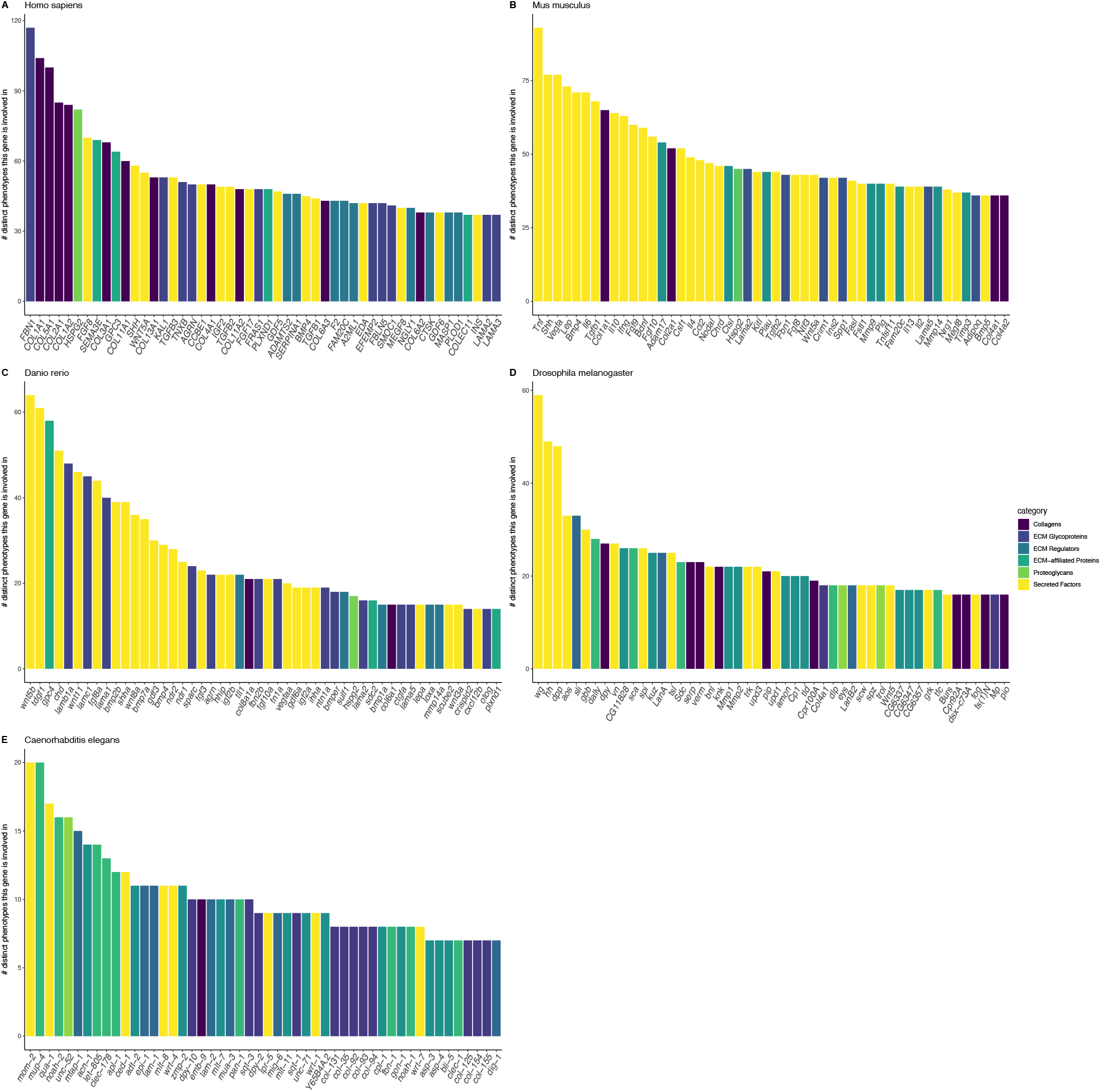
Degree of phenotypic impact of matrisome genes (grouped) The number of phenotypes that associate with each matrisome gene are shown in decreasing order for the top 50 genes. This graph is complementary to Fig. 3., to show the impact of phenotypic grouping on the number of phenotypes. The gene name is displayed on the x-axis and the number of unique phenotypes on the y-axis. The color of the bars refers to the matrisome category for each gene.

**Supplementary Fig. 6:**
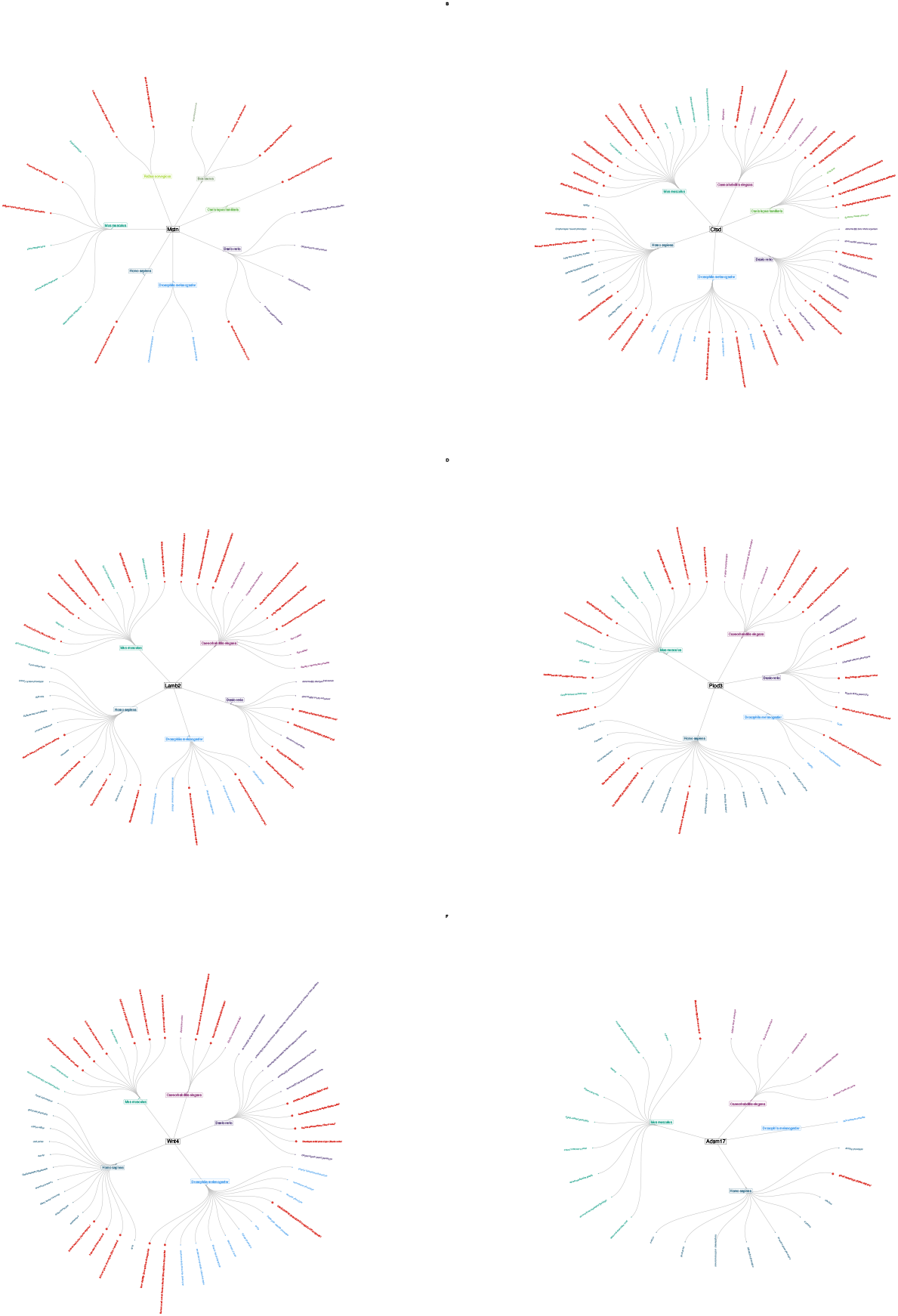
Phenotypic fingerprint of the most conserved matrisome members. Circular dendrograms depicting the cross-species phenotypic fingerprint of all human matrisome genes for which at least one phenotype has been studied in at least one other species. The human orthologue is shown in the middle connected to the species in which the gene was associated with a phenotype, and then each connected to terminal nodes depicting the grouped phenotypes. Phenotypes that were observed in more than one species are highlighted in red and the size of the terminal node reflects the number of species this phenotype was observed in.

**Supplementary Fig. 7:**
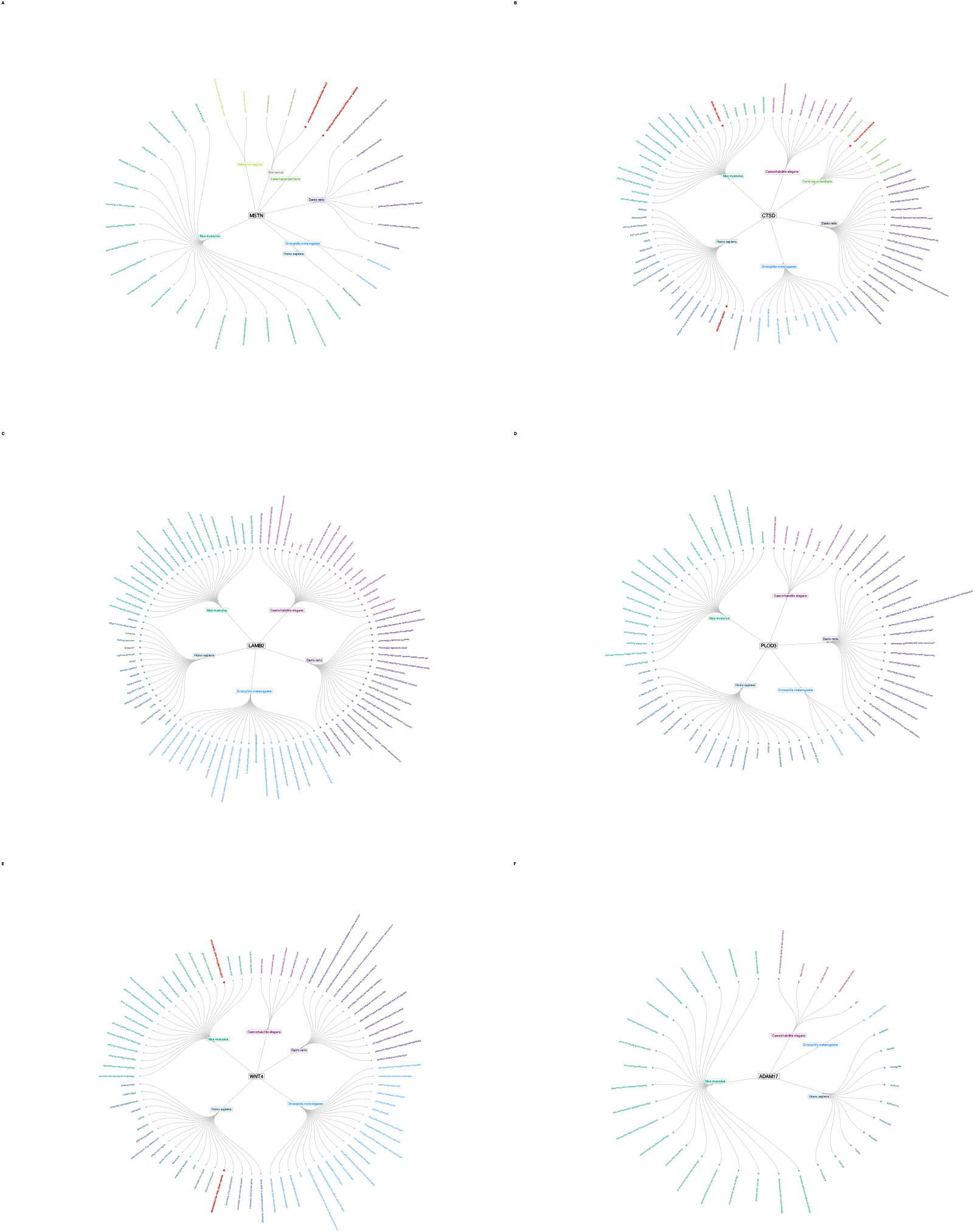
Phenotypic fingerprint of the most conserved matrisome members (ungrouped) Circular dendrograms depicting the cross-species phenotypic fingerprint of all human matrisome genes for which at least one phenotype has been studied in at least one other species. This figure is complementary to Supplementary Fig. 6, but in contrast is showing the phenotypic fingerprint for ungrouped phenotypes.

**Supplementary Fig. 8:**
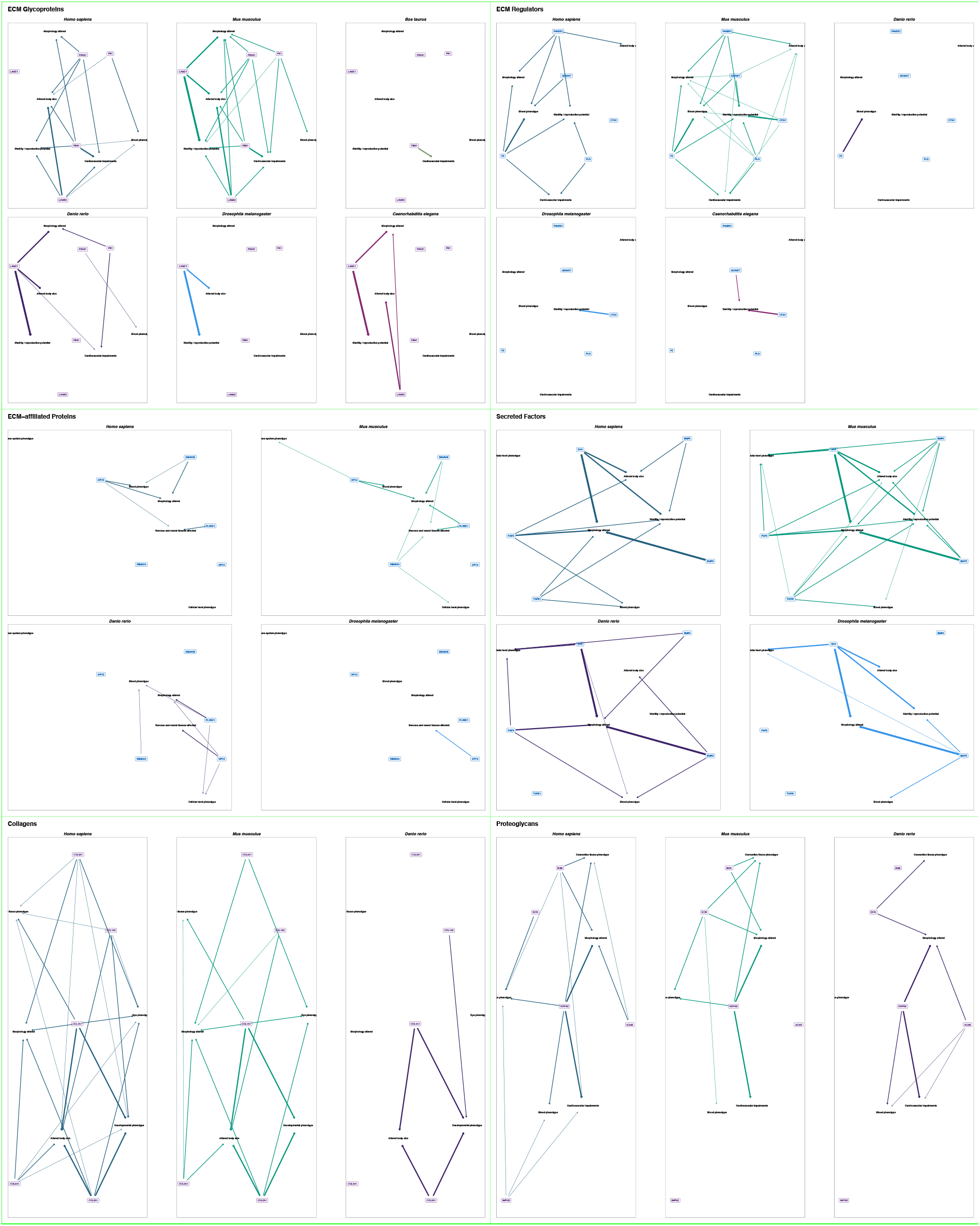
Phenotypic implications of the matrisome categories across species. For each matrisome category, the five genes and phenotypes with the highest connectivity across species are highlighted for each taxon. Grouped phenotypes are shown as plain text while genes are shown as colored labels always referring to their human orthologues. If a gene has been associated with at least one phenotype associated with the phenotype group they are connected with an arrow. The degree of conservation of the gene-to-phenotype association is reflected by the width and opacity of the connecting arrow.

**Supplementary Fig. 9:**
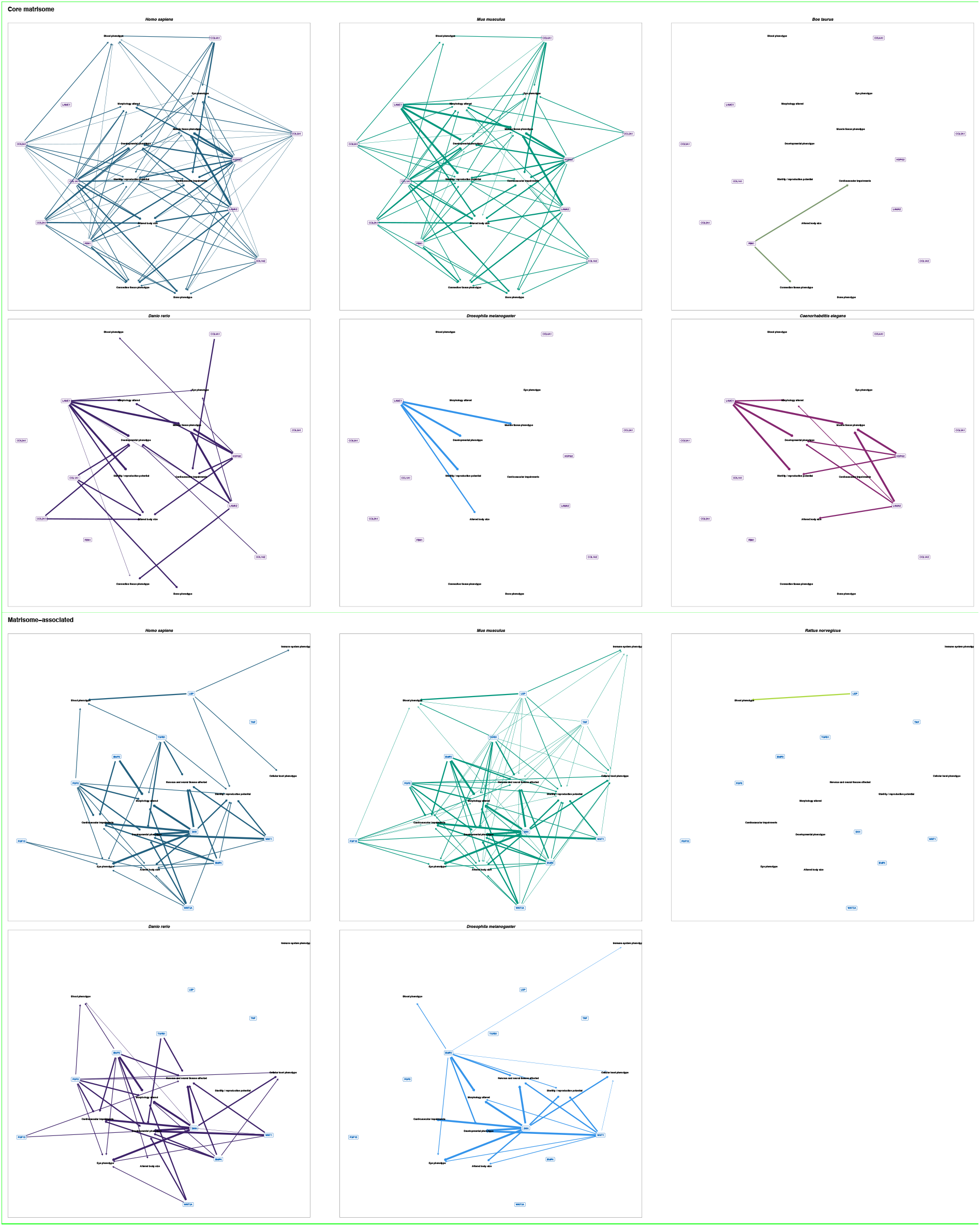
Phenotypic implications of the matrisome divisions across species. For each matrisome division, the ten genes and phenotypes with the highest connectivity across species are highlighted for each taxon. Grouped phenotypes are shown as plain text while genes are shown as colored labels always referring to their human orthologues. If a gene has been associated with at least one phenotype associated with the phenotype group they are connected with an arrow. The degree of conservation of the gene-to-phenotype association is reflected by the width and opacity of the connecting arrow.

## Discussion

The recent collective scientific efforts focusing on deep phenotyping and phenome-wide association studies revealed many novel genotype-to-phenotype relationships [2,3,29]. Extracellular matrices are important for cellular and organismal function [13,17]. However, the contribution of ECM gene variants to the phenotypic landscape is unknown.

Here, we defined the ECM phenome of humans, mice, zebrafish, *Drosophila*, and *C. elegans*, with extrapolations to three other species. We found over 42’558 matrisome genotype-to-phenotype relationships across the five defined matrisome species and an additional 306 genotype-to-phenotype relationships in the three undefined-matrisome species. We identified the phenotypic landscape of the matrisome that is conserved among species and also species-specific phenomes. Our analysis of the phenotypic fingerprints of genes revealed *MSTN, CTSD, LAMB2, HSPG2*, and *COL11A2* matrisome genes bearing the most associated phenotypes. From these genotype-to-phenotype interactions, we have built networks linking analogue phenotypes that might be driven by similar underlying molecular mechanisms involving matrisome genes. We provide the ECM phenome as a platform to use information gained from model organisms to implicate novel genotype-to-phenotype relationships for humans.

One limitation of our analysis are our nearly-completed phenotype categorical collections, which include the most frequent occurring phenotypes, but ignores some infrequently occurring phenotypes during our manual curation process due to the large number phenotypes and sometimes jargon-specific phenotype naming. We aimed to functionally group phenotypes so that these become more comparable across species. This includes also species-specific phenotypes. We manually added 156 broadly defined phenotype categories, which captured 86.8 % of the original phenome. However, while the grouping achieves its primary goal to unify the most frequent phenotypes some of the less abundant phenotypes might be falsely grouped or remain ungrouped. Modifications in the grouping system achieved a better assignment of misclassified infrequent phenotypes, with little or no changes in the top phenotypic categories, which adds reliability to our findings and interpretations in this present study. All analysis including grouped or ungrouped are provided in the supplementary sections, allowing researchers to build upon.

The strong points in the present study are our networks of matrisome genes mapped to recorded phenotypes, which offers a conceptual framework for further investigation. This phenome-based network allows the identification of conserved underlying molecular interactions of matrisome genes across comparable phenotypes and species. In particular, it enables queries on given human phenotypes or ECM genes, which are provided with genotype-to-phenotype relationships across other species, suggesting potential novel molecular targets or read-outs for phenotypic high-throughput screens. For instance, early developmental or lethal genes that lead to spontaneous abortion in humans, can be studied with their orthologue gene counterparts in other species [10]. We provide the most conserved subset of the matrisome-interactome dataset in graphical format highlighting the genes and phenotypes with the highest degree of connectivity (Fig. 5 and 6; Supplementary Fig. 8 and 9). Such graphical interaction frameworks have been invaluable for all known human gene associated phenotypes [10].

In our phenome analysis across species, we identified novel aging-related phenotypes associated with matrisome gene variants in the top categories (Fig. 2). Decline and remodeling of the matrisome is a major driver of aging and longevity [18,30–32]. Recently, the human aging phenome has been defined revealing that phenotypes associated with collagens or matrisome, such as facial wrinkles, kyphosis, arthritis, and osteoporosis are under the most prevalent phenotypes recorded from 77 million elderly [8]. Furthermore, connective tissue diseases, such as Marfan syndrome caused by mutation in *FBN1* and Ehlers-Danlos syndrome, were clustered with features of premature aging diseases (progeria) and aging [8]. In our analysis, we found that the *FBN1* gene has the most associated human phenotypes (Fig. 3 and Supplementary Fig. 5).

The breath of the phenotypic fingerprints of some matrisome genes hints at the conserved modularity of gene systems. Some of these orthologue phenotypes across species suggest a repurposing of the underlying molecular systems through evolution. Our network of these matrisome genes associated with these orthologues phenotypes facilitate studying human phenotypes or even diseases in nonobvious model systems. Thus, our computational analysis provides a systems-level approach to facilitate mechanistic discoveries underlying ECM phenomes across species.

## Author contributions

All authors participated in analyzing and interpreting the data. CYE and CS designed the computational analysis and wrote the manuscript.

## Author Information

The authors have no competing interests to declare. Correspondence should be addressed to C. Y. E.

## Acknowledgement

We thank members of the Ewald lab for critical reading of the manuscript, Davide Vitiello for his help with the phenotypic grouping, and the Monarch Initiative (https://monarchinitiative.org) for providing the indispensable resources for the genotype-phenotype analysis performed in this work. Funding for this study was from the Swiss National Science Foundation grant number PP00P3_163898 to CS and CYE.

